# Managing populations after a disease outbreak: exploration of epidemiological consequences of managed host reintroduction following disease-driven host decline

**DOI:** 10.1101/2025.02.28.640833

**Authors:** Jorge Arroyo-Esquivel, Alyssa Gehman, Ken Collins, Fabio Sanchez

**Affiliations:** California Department of Fish and Wildlife, California, USA; Hakai Institute, British Columbia, Canada; University of British Columbia, British Columbia, Canada; Unaffiliated researcher; Escuela de Matemática, Universidad de Costa Rica. San José, Costa Rica; Centro de Investigación en Matemática Pura y Aplicada. San José, Costa Rica

**Keywords:** disease-driven mortality, *Pycnopodia helianthioides*, sea star wasting disease, reintroduction, age-stage model

## Abstract

Disease outbreaks in wild populations around the globe can lead to widespread mortality within populations, where recovery of individuals can be rare. An example of this population is the sunflower sea star *Pyc-nopodia helianthioides* in the Northeastern Pacific coast. The populations of this species, as well as many other sea star populations, have experienced massive mortality events due to an unidentified disease called Sea Star Wasting Disease (SSWD).

*Pycnopodia* play a key role in providing top-down control of kelp grazers in rocky reefs across the Northeastern Pacific coast. This, combined with the massive declines in kelp coverage observed during the 2015-2016 marine heat wave observed in the Northeastern Pacific, has sparked an interest in reintroducing *Pycnopodia* individuals in the coast to potentially assist in recovery of the populations. However, the epidemiological implications of reintroducing healthy sea stars into the wild populations is an under-explored question. This work explores this question using a dynamical population model of *Pycnopodia*. We use this model to estimate the impacts of reintroducing healthy individuals into a wild sea star population. This analysis will provide valuable information for managers interested in restoring *Pycnopodia* populations.

## Introduction

Disease-induced mortality can lead to significant decreases in the density of natural populations, increasing their risk of extinction [23]. One of the most studied examples is the mass mortality of amphibian populations worldwide caused by the fungal pathogen *Batrachochytrium dendrobatidis* [5, 19]. Effectively managing a population heavily affected by disease-induced mortality has several challenges and depends on the nature of transmission (frequency vs. density dependent), infected individuals’ recovery rates, and the population density of both infected and total individuals [1].

A possible management strategy for populations affected by disease-induced mortality is reintroducing healthy individuals into the population, either from other natural populations or conservation breeding facilities [7]. However, this strategy presents several risks, particularly if the disease has not been eradicated in the population. Introducing new individuals may affect genetic diversity and cause outbreeding depression in the population [10]. In addition, an increase in host density can lead to an increase in disease prevalence, which would cause the contrary effect to the actual goal [18].

An example of a population of conservation interest where the reintroduction of healthy individuals has been considered is the sunflower sea star, *Pycnopodia helianthioides* [12, 15]. This sea star has been heavily affected by the sea star wasting disease (SSWD). This, in addition to extreme marine heatwaves, has caused many populations of this sea star in the Northeastern Pacific to experience substantial declines, even becoming functionally extinct in some areas such as the northern coast of California [14]. Particular interest in the recovery of sunflower sea star populations has been driven by its role as a predator of purple sea urchins (*Strongylocentrotus purpuratus*). The loss of top-down control of purple sea urchins by sunflower sea stars has been identified as potentially one of the main drivers of kelp decline in the northern coast of California [24]. Furthermore, the recovery of the sunflower population is potentially key to a more resilient kelp forest after recovery [11, 2].

Given the ecological impacts of the sunflower sea star population’s demise, there is a strong interest in reintroducing individuals to the regions where the population has become functionally extinct. Although significant progress has been made on the technical methodology of breeding individual sea stars [16], no evaluation of the potential disease impacts of these reintroductions has been done. Evaluation of the potential disease impacts can enable managers to take steps to avoid potential pitfalls and hopefully provide the best possible chance of successful recovery.

In this paper, we evaluate the population effects of introducing healthy sea stars in a population using a simple age-stage model of the sunflower sea star population. The following section details the model and how we analyze it. Following it, we present how the population growth rate is affected by different introduction strategies, both from an ecological and an eco-evolutionary perspective. We finish this paper by discussing how these results can inform managers in future sunflower sea star reintroduction efforts.

## 2 Methods

### 2.1 Modeling

Our age-stage model follows the abundance of a sea star population separated into three classes: juveniles *J*, adults *A*, and infected *I* individuals. The model follows these abundances through seasons *t*, with *t* = 0 representing winter. A season-dependent number of juveniles *r*_*t*_ are produced for the following season. Therefore, we assume that recruitment occurs constantly throughout the year. This coincides with the observations in both of our sites where recruits were observed throughout the year, as well as with classic studies that reported spawning seasons of December-January and March-July, with metamorphosis of larvae occurring anywhere between 90 and 146 days post spawning [25]. A fraction *α* of juveniles mature into adults. A proportion *µ*_*S*_ of all healthy juveniles and adults experience mortality at a season.

For infected individuals, a season-dependent proportion *β*_*t*_ of individuals become infected. Due to the possibility of sea stars of different populations infecting our model population [17], we assume this infection is frequency-dependent. Recovery of *Pycnopodia* from SSWD is rare, and thus, we assume that an infected sea star cannot return to a healthy state [4]. Finally, a proportion *µ*_*I*_ of those infected individuals experience mortality at a season.

We incorporate two additional control parameters to model the effects of introducing healthy individuals. These parameters correspond to the introduction of healthy juveniles *λ*_*J*_ and adults *λ*_*A*_.

In equation form, our model follows:

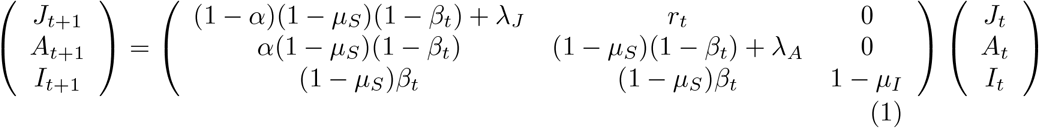

### 2.2 Parameter estimation

We estimate our model parameters (Table 1) using recruitment surveys from two regions in the northeastern Pacific coast: Saratoga Beach (Island County, WA, USA) and the central coast of British Columbia, Canada.

**Table 1:** Parameters of Equation 1 and their baseline values, including their range of values explored in the global sensitivity analysis.

| Parameter | Description | Baseline Value | Range |
| --- | --- | --- | --- |
| $r_t$ | Baseline recruitment proportion of juvenile sea stars | Winter: 0.55 recruits/adult<br>Spring: 0.71 recruits/adult<br>Summer: 0.59 recruits/adult<br>Fall: 0.35 recruits/adult | [0, 1] |
| $\beta_t$ | Baseline infection proportion of sea stars | Not fall: 0<br>Fall: 0.2 | [0, 1] |
| $\alpha$ | Baseline maturation proportion of sea stars | 0.18 | [0, 0.5] |
| $\mu_S$ | Mortality proportion of healthy sea stars | 0.005 | [0, 0.05] |
| $\mu_I$ | Mortality proportion of infected sea stars | 0.95 | [0.9, 1] |

The study site at Saratoga Beach is a sandy environment located at the seaward edge of an extensive eelgrass (*Zostera marina*) bed. Two sea star species (*Pisaster brevispinus, Pycnopodia helianthoides*) are present and a third (*Luidia foliolata*) is rare. Since Spring of 2020, monthly surveys have been conducted along depth contours in four sub-habitats; in and on the edge of the eelgrass bed, in an area of seasonal kelp (*Saccharina latissima*), and open sand. Depth ranges from 4-10 m (MLLW). Four 10 *×* 2*m* transects are surveyed in each sub-habitat, for a total site area of 320 *m*^2^. Every sea star is counted, measured, and assessed for SSWD. Additionally, the associated substrate and activity level for each *Pycnopodia* is recorded.

The study site in British Columbia includes 165 sites, conducted from 2013-2024 at a range of habitat types, including rocky reefs, seagrass habitats, soft sediments, and kelp forests (Gehman et. al, Proceedings B, accepted). At each site, 30×2m transects were surveyed, with 3-8 surveys per sites. Sites are surveyed in May, July, August and October (although not evenly at all sites or all years). Depth range was between 0-10m (Canadian Chart Datum). Individuals were considered recruits when they were *<* 2*cm* radius.

We calculate the recruitment proportion at each season *r*_*t*_ by dividing the mean number of recruits observed in these sites at a given season by the number of adults observed. The infection proportion *β* is calculated using line search by running the model with different values of *β* in order for the model to match the mean proportion of *Pycnopodia* detected with SSWD (approximately 15% for both sites). Baseline maturation *α* was determined from the growth rate of juveniles calculated by [21] at approximately 8cm/year. We then calculate *α* using this growth rate and the assumption that *Pycnopodia* reaches sexual maturity at a diameter of 11cm (Sarah Gravem, personal communication). Mortality proportion of healthy and infected individuals are assumed to be extremely low and high, respectively.

### 2.3 Data and model analysis

We analyzed how a *Pycnopodia* population is affected by reintroduction efforts by estimating the population growth rate of Model 1 using the baseline values provided in Table 1. We start our simulations with an arbitrary initial population abundance *N*_0_ where 25% of the population consists of juveniles, 74.25% consists of adults, and 0.75% consists of infected individuals. Because the parameters of this model vary through time, we estimate the population growth rate *λ* using the geometric mean through time *GM*_*t*_ of the ratios of the season-over-season population abundances [20]. In equation form, this corresponds to:

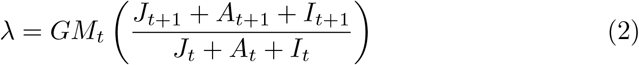

We vary reintroduction efforts by timing (in what season reintroduction occurs) and intensity. We vary the intensity of reintroduction relative to the current abundance, i.e., an intensity of 10% corresponds to reintroducing several individuals equal to 10% of the number of individuals of the corresponding age. To evaluate the possibility of reintroduced individuals positively affecting the resistance to the disease, we test two scenarios, one where the proportion of individuals infected *β*_*t*_ is fixed, and one where this proportion is reduced due to restoration efforts.

We perform a Global Sensitivity Analysis (GSA) following the methods described in [13] to understand the relative importance of the different model parameters in the population growth rate. The most critical parameters that determine the growth rate are identified using the importance metric of a random forest analysis, where a combination of different parameter values is used to predict the population growth rate. This metric is a relative measure of how much varying an individual parameter leads to a variation in the growth rate predicted by the trained random forest. We sample parameters from uniform distributions following the range for each parameter presented in Table 1. In addition, we vary the intensity of reintroduction in the range of 0 − 50% intensity, with 10% of parameter combinations having no reintroduction intensity.

## 3 Results

Our model findings provide critical insights into the reintroduction dynamics of *Pycnopodia* populations, revealing key interactions between infection rates, reintroduction strategies, and long-term population viability. The results underscore the delicate balance between population growth and disease prevalence, demonstrating the reintroduction efforts may fail to sustain viable populations without careful management. By analyzing various scenarios, we identify strategic approaches that can maximize the effectiveness of conservation interventions. The relationship between population growth rate (*λ*) and infection rate (*β*) reveals a strong inverse correlation. As infection rates rise, population growth declines, and a critical threshold emerges where even small increases in *β* can drive long-term population decline. Figure 1 illustrates this trend, emphasizing the importance of disease management as a fundamental component of reintroduction efforts.

**Figure 1:**
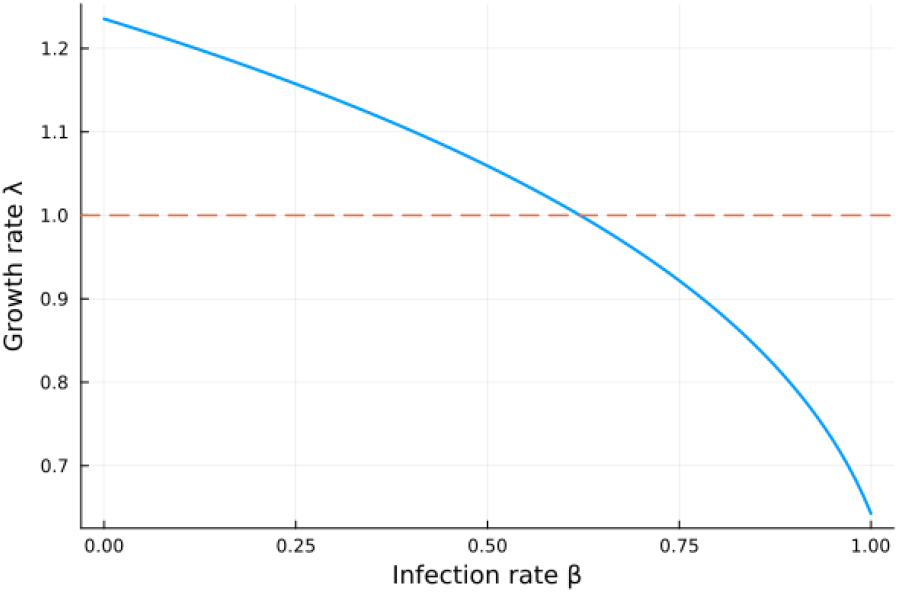
Population growth rate *λ* of Model 1 as the proportion of infected individuals *β* varies. The horizontal line corresponds to *λ* = 1, which separates the parameter values whether a population grows (*λ >* 1) or decreases (*λ <* 1) in the long term.

Moreover, our model suggests seasonal timing plays a crucial role in determining the success of reintroduction efforts. Modeled reintroductions conducted in spring and summer yield the highest population growth rates, particularly when juvenile reintroduction exceeds 30% of the current population (Figure 2). This provides a framework for timing interventions to coincide with these favorable recruitment conditions that can enhance reintroduction success. Furthermore, our model indicates that while moderate-intensity reintroductions (20% − 30%) contribute significantly to population stability, there are diminishing returns beyond a certain threshold, emphasizing the importance of optimizing effort allocation rather than maximizing intensity.

**Figure 2:**
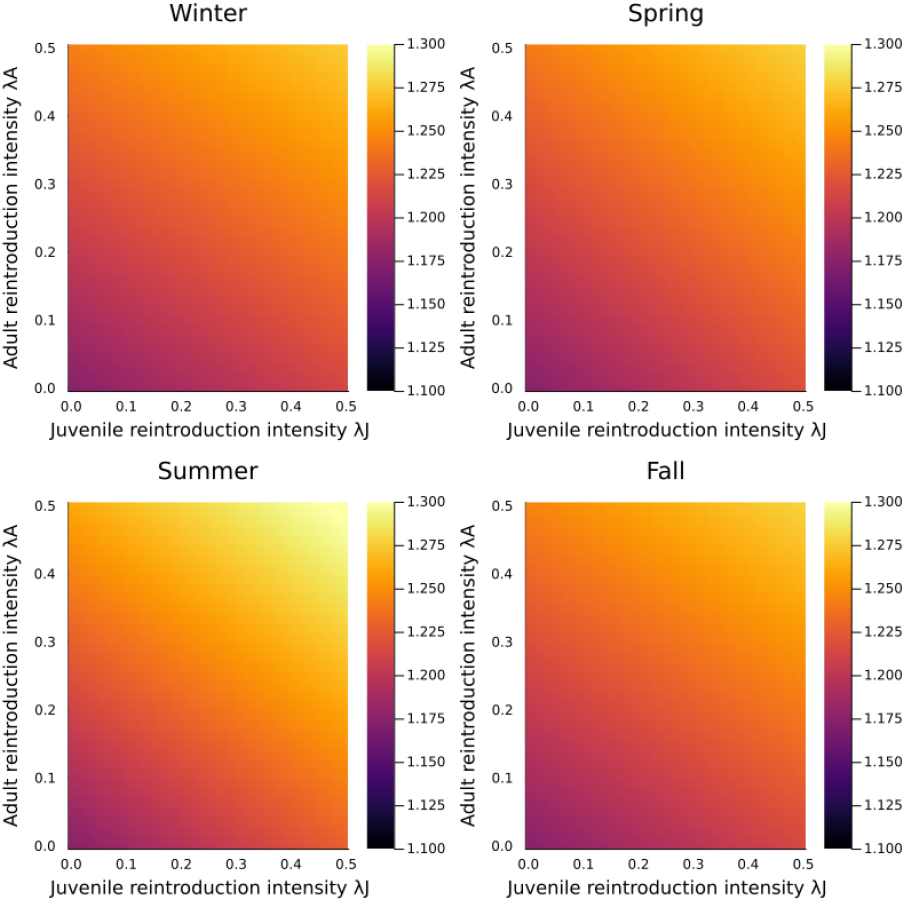
Population growth rate *λ* of Model 1 at different reintroduction intensities (as percent of total abundance) of juvenile and adult individuals at different seasons.

Reintroduction duration further influences population recovery. Our model results demonstrate that sustained efforts over multiple seasons lead to significantly higher long-term growth rates than short-term interventions. Even moderate reintroduction levels contribute to enhanced population resilience when maintained across several seasons. This effect is particularly pronounced when reintroduction efforts reduce infection prevalence, as newly introduced healthy individuals dilute the proportion of infected individuals within the population. Figures 3 and 4 illustrate this dual effect, direct demographic support and epidemiological mitigation, underscoring the necessity of long-term commitment to reintroduction programs.

**Figure 3:**
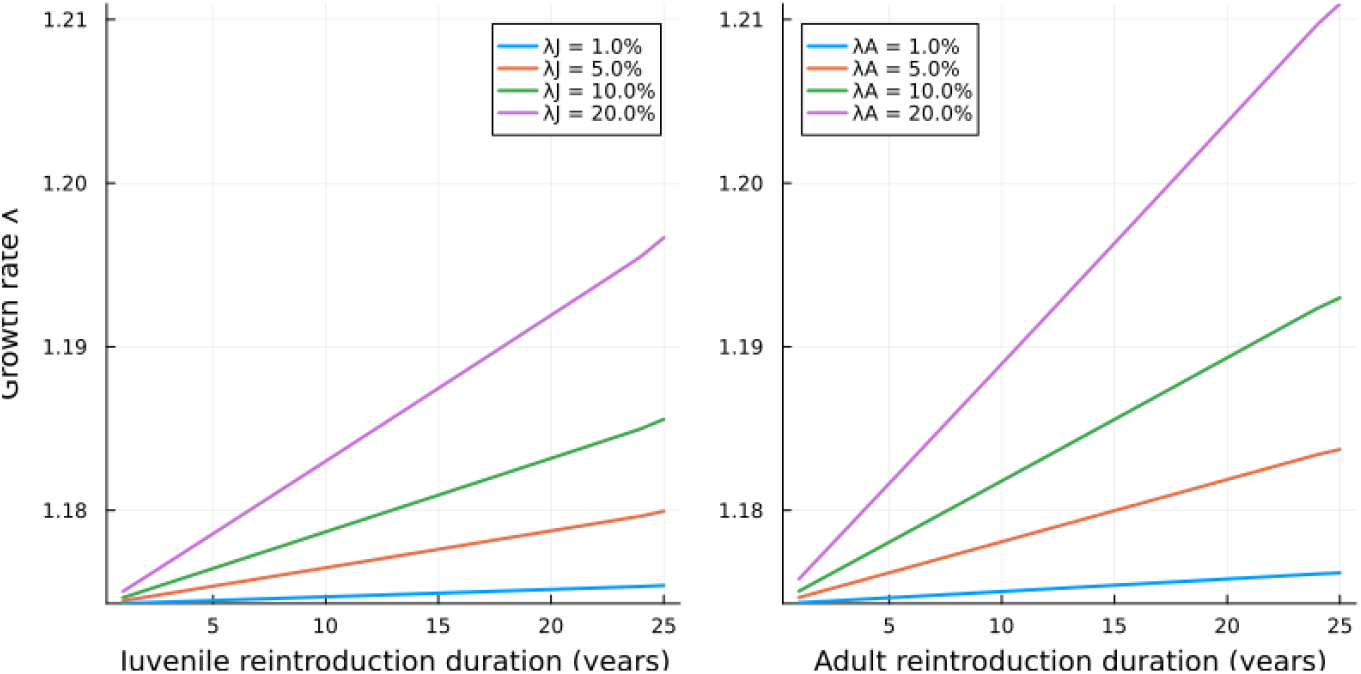
Population growth rate *λ* of Model 1 at different summer reintroduction intensities (as percent of total abundance) and durations if the proportion of infected individuals *β* does not change with the introduction of new individuals.

**Figure 4:**
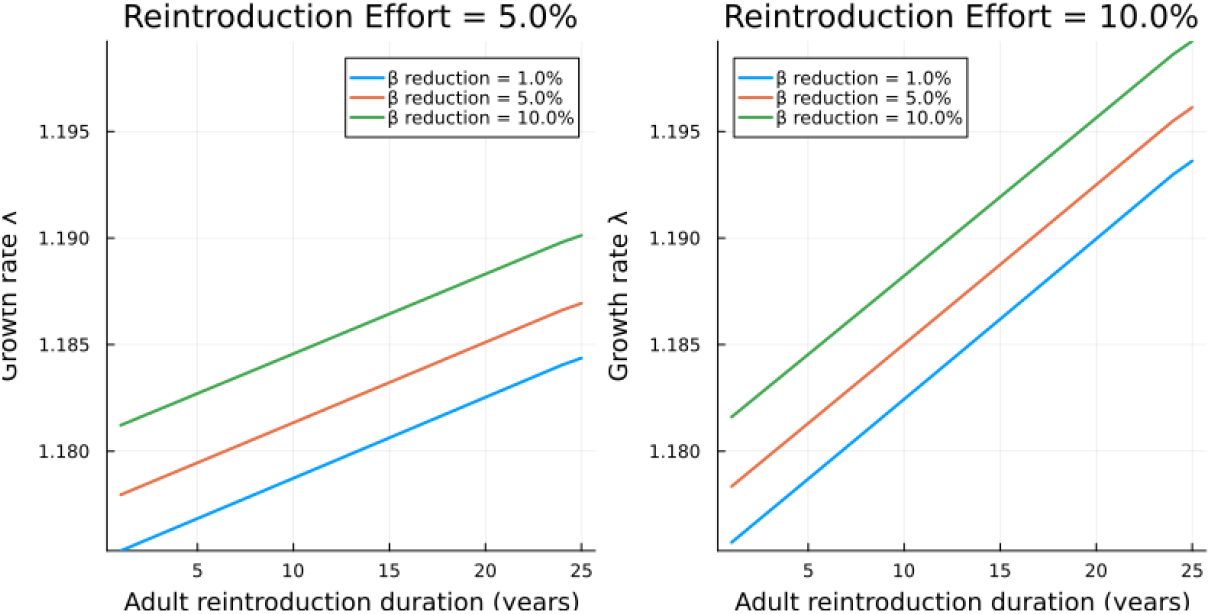
Population growth rate *λ* of Model 1 at different summer reintroduction intensities (as percent of total abundance) and durations in the case the proportion of infected individuals *β* changes with the introduction of new individuals.

Global sensitivity analysis (GSA) confirms that infection rate (*β*) is the most influential factor affecting population growth in our model. This reinforces the need for targeted disease management alongside reintroduction efforts. Additionally, the maturation rate (*α*) and juvenile recruitment rate (*r*_*t*_) significantly impact *λ*, suggesting that promoting natural recruitment through habitat restoration or improved larval survival conditions can further enhance recovery efforts. Figures 5 and 6 highlight the relative influence of key model parameters, with infection prevalence as the primary driver of population trends. The high impact of *β* in Figure 5 suggests that in the absence of a decrease in infection prevalence, the reintroduction success remains limited. Conversely, Figure 6 shows that when infection prevalence is mitigated through management strategies, reintroduction becomes far more effective.

**Figure 5:**
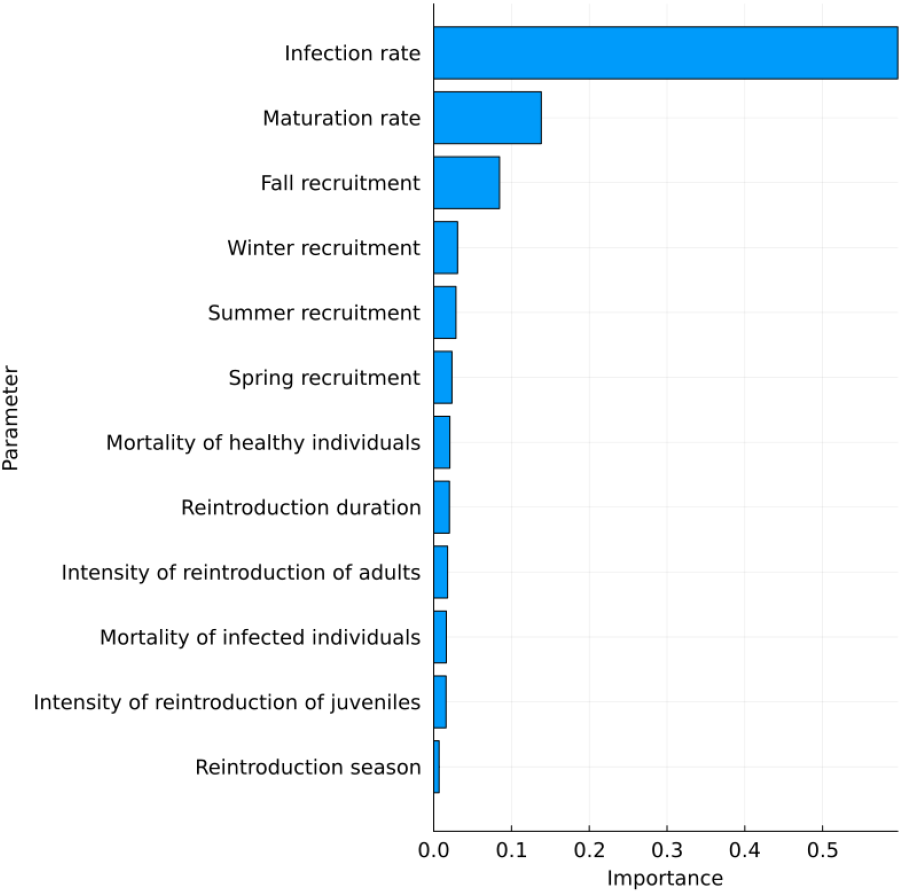
Parameter importance of the different parameters to estimate the long-term growth rate *λ* of Model 1 determined by the GSA in the case the proportion of infected individuals *β* does not change with the introduction of new individuals.

**Figure 6:**
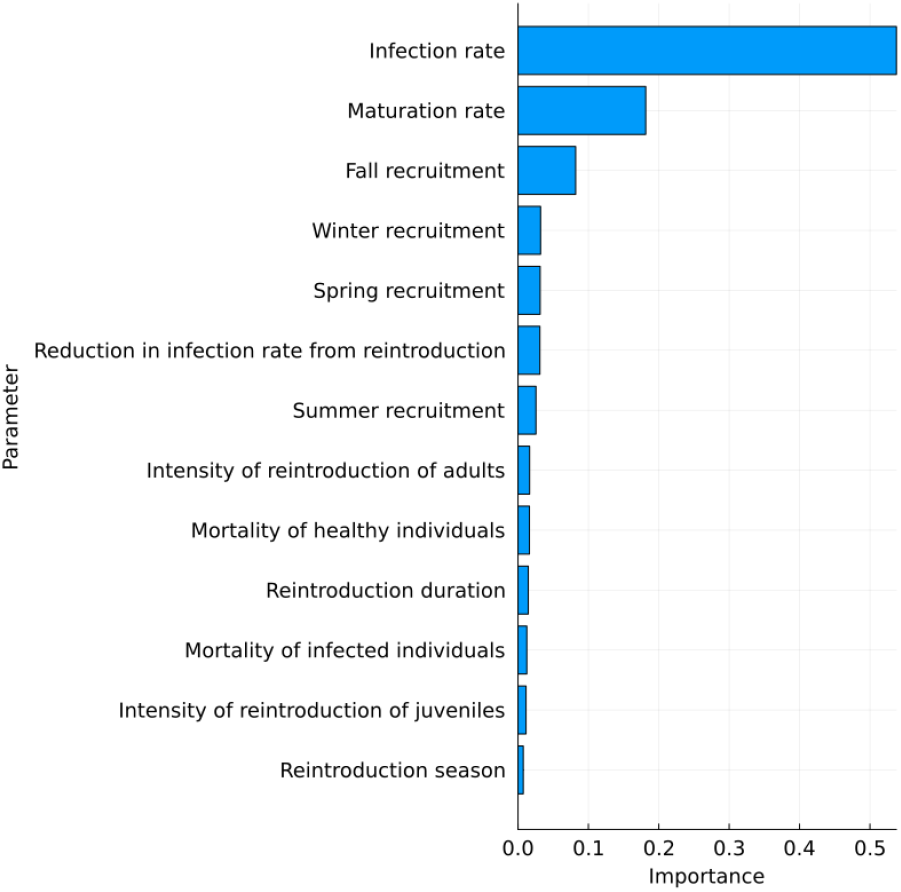
Parameter importance of the different parameters to estimate the long-term growth rate *λ* of Model 1 determined by the GSA in the case the proportion of infected individuals *β* changes with the introduction of new individuals.

Our model suggests targeted seasonal reintroductions in spring and summer have the potential to maximize recruitment success and resilience. Long-term reintroduction commitments yield better outcomes than short-term interventions, reinforcing the importance of sustained efforts. Disease management is crucial to success, as unchecked infection rates can undermine even the most aggressive reintroduction programs.

## 4 Discussion

This study identifies the primary drivers of population growth in a two-age-class model population affected by a disease without recovery. Our findings indicate that the disease’s infection rate predominantly influences the population growth rate. This result aligns with previous analyses, suggesting that high infection rates in a system with no recovery can prevent population persistence without external inputs [4]. Additionally, maturation rate exerts a smaller yet significant effect on population dynamics, consistent with findings in other marine organisms [6].

In environments where the population can persist despite disease pressure, our results suggest that reintroduction efforts have a limited to moderate impact on population growth. However, our model indicates that this impact can be amplified by reintroducing adult individuals, particularly during the summer. A possible explanation for this outcome is that new adults introduced in summer may compensate for those lost to infection in the subsequent fall. While our model suggests that adult reintroduction exerts the most significant effect on population growth, empirical studies indicate that introducing individuals across multiple age classes can enhance the success of reintroduction efforts [18].

The relatively low sensitivity of population growth to reintroduction intensity suggests that these results may be generalizable to sites where infection rates are high enough to prevent long-term persistence. In such cases, reintroduction efforts are likely most effective in areas where *Pycnopodia* has become functionally extinct and where the reintroduced individuals exhibit genetic resistance to the pathogen. Thus, breeding for genetic resistance for reintroduced individuals could expand the range of conditions wherein reintroduction efforts are successful.

Another key environmental factor influencing the population dynamics of *Pycnopodia* and other marine invertebrates is El Niño. The decline of *Pycnopodia* and other sea star populations over the past decade has been attributed to a combination of an intense El Niño event and a surge in sea star wasting disease (SSWD) [22]. Previous studies have linked these SSWD outbreaks to the elevated sea temperatures associated with El Niño and the Blob of 1984 [8, 3, 9]. It is plausible that infection rates temporarily increase during these climatic events, leading to mass mortality and a subsequent decline in infection rates once the outbreak subsides. However, the precise mechanisms through which El Niño affects marine invertebrate population dynamics remain poorly understood. If temperature and disease are interacting, then heatwave conditions could potentially push populations to high enough disease transmission levels to keep the population growth rate low, despite high levels of introduction efforts previous to a heatwave. Improved knowledge of these interactions would facilitate the development of more accurate models for managing populations under El Niño conditions.

Beyond the omission of El Niño effects, our model also assumes that *Pycnopodia* recruitment occurs consistently each year and across all seasons. While no direct evidence contradicts this assumption, data from Saratoga Beach, WA indicate four recruitment events over the past six years, suggesting that recruitment may not occur yearly. The underlying drivers of this variability remain unknown, complicating efforts to incorporate irregular spawning into population models. If recruitment is episodic rather than continuous, we expect a reduction in population growth rates, potentially leading to population decline in specific sites. Under such conditions, the impact of reintroduction efforts would likely increase, as introducing healthy individuals could help compensate for years or seasons with limited natural recruitment.

This study highlights the complexity of post-disease population recovery and underscores the necessity of integrating epidemiological considerations into conservation planning. In addition, the model introduced in this paper has the potential to be used or extended to assist in the understanding of disease dynamics for *Pycnopodia* and other sea star populations affected by SSWD. Future research should incorporate additional ecological drivers, such as climate variability and genetic resistance, to refine and optimize reintroduction strategies.

